# A reversible temperature block captures pre-lysosomal intermediates of starvation-induced ERphagy

**DOI:** 10.64898/2026.10.06.756943

**Authors:** Marco Fabbro, Tatiana Soldà, Diego Morone, Carmela Galli, Andrea Raimondi, Maurizio Molinari

## Abstract

The selective lysosomal turnover of endoplasmic reticulum (ER) portions, termed ERphagy, removes aged, excess or damaged ER domains. The intermediates forming between receptor activation and lysosomal delivery are short-lived and remain poorly resolved. Here we identify a reversible temperature block at 25°C that arrests ERphagy before lysosomal delivery. At this temperature, nutrient restriction still drives focal accumulation of the endogenously Halo-tagged ERphagy receptor TEX264 (puncta formation), segregation of ER cargo, remodelling of the ER membrane, whereas delivery of this material to degradative endolysosomes is strongly reduced. Rewarming above 27°C restores lysosomal delivery, showing that the accumulated eTEX264-Halo puncta correspond to pre-lysosomal intermediates of the ERphagy program. Their formation requires ULK1/2 activity, the ATG7-dependent conjugation machinery, ATG9A, ATG13 and FIP200, coupling receptor clustering to autophagosome biogenesis. Consistently, expansion microscopy and electron tomography reveal curved, cradle-like TEX264-positive ER domains containing ER cargo, co-localizing with autophagy markers and lying adjacent to p62-cargo condensates.

## Introduction

The lysosomal turnover of the ER, ERphagy, is mediated by a set of membrane-associated receptors, including the FAM134 variants (Khaminets *et al*, 2015), SEC62 (Fumagalli *et al*, 2016), RTN3L (Grumati *et al*, 2017), CCPG1 (Smith *et al*, 2018), and TEX264 (An *et al*, 2019; Chino *et al*, 2019). These receptors occupy distinct ER subdomains and, in some cases, exhibit tissue-specific expression patterns enabling selective remodelling of the ER in response to different demands and stimuli including cell differentiation, nutrient deprivation, ER stress, and the accumulation of misfolded proteins (Chino & Mizushima, 2023; Gubas & Dikic, 2022; Mochida & Nakatogawa, 2022; Reggiori & Molinari, 2022). TEX264 is a major ERphagy receptor during nutrient restriction and contributes substantially to starvation-induced ER turnover (An *et al*., 2019; Chino *et al*., 2019). Upon nutrient restriction, TEX264 transiently concentrates in discrete puncta, frequently at ER tubular three-way junctions before being rapidly delivered, together with ER material, to degradative endolysosomes (An *et al*., 2019; Chino *et al*., 2019). Studies from several laboratories have shown that increased expression of fluorescently tagged ERphagy receptors can promote their post translational modifications and clustering, ER remodelling and lysosomal delivery of ER material, i.e., it phenocopies ERphagy induction (e.g., (Foronda *et al*, 2023; Gonzalez *et al*, 2023)). Although these approaches have been instrumental in defining receptor function, ERphagy receptors overexpression bypasses regulatory events controlling receptor abundance, availability, activation and assembly, and provides limited information on the behavior of endogenous receptors during the physiologic initiation of ERphagy (An *et al*., 2019; Chino *et al*., 2019; Fumagalli *et al*., 2016; Grumati *et al*., 2017; Khaminets *et al*., 2015; Liang *et al*, 2018; Rudinskiy *et al*, 2025; Rudinskiy *et al*, 2024; Smith *et al*., 2018). To overcome these limitations and with a particular interest in monitoring the phases of starvation-induced ERphagy that precede lysosomal delivery under endogenous receptor expression, we built on previous work showing that C-terminal fusion of TEX264 does not impair its distribution within the ER and its function as an ERphagy receptor (An *et al*., 2019; Chino *et al*., 2019; Kucinska *et al*, 2023) and generated mouse embryonic fibroblasts (MEFs), where the endogenous (e)TEX264 locus was C-terminally fused to HaloTag by CRISPR/Cas9 genome editing (eTEX264-Halo). HaloTag is a 33kDa protein tag derived from a bacterial haloalkane dehalogenase that covalently binds chloroalkane-conjugated ligands (Los *et al*, 2008), enabling visualization of tagged proteins by fluorescence and electron microscopy, biochemical analyses and live cell imaging (Fregno *et al*, 2018; Rudinskiy *et al*, 2022; Yim *et al*, 2022). Live-cell studies of ATG protein recruitment indicate that individual autophagosome formation events are completed within minutes (Koyama-Honda *et al*, 2013), so that at any given time only a few nascent structures are present per cell and their membrane architecture and molecular organization is poorly defined, if at all. Strategies to overcome this limitation have so far relied on genetic or pharmacological interference with the pathway, such as depletion of ESCRT (Loi *et al*, 2019; Schafer *et al*, 2020; Takahashi *et al*, 2018), or contact sites components (Fasana *et al*, 2025), or ultrastructural analyses of autophagy-deficient cells (Kishi-Itakura *et al*, 2014). These approaches enrich intermediates but arrest them irreversibly, precluding any test of whether the accumulated structures remain competent for endolysosomal delivery.

Here, by monitoring starvation-induced ERphagy in cells expressing endogenous eTEX264-Halo, we show that nutrient restriction at 25°C drives focal accumulation of endogenously Halo-tagged TEX264, spatial segregation of ER cargo, remodelling of the ER membrane but the pathway stalls before TEX264-positive ER material reaches the endolysosomes. This temperature-sensitive block is reversible and returning starved cells to 37°C restored delivery of eTEX264-Halo positive ER material to degradative endolysosomes. Live-cell imaging in which the temperature of the culture medium was recorded throughout, indicated that delivery can resume above the 27°C, when individual eTEX264-Halo puncta were observed entering the degradative compartments. Exploiting this conditional arrest together with confocal laser scanning microscopy (CLSM), expansion microscopy (ExM) and correlative light and electron microscopy (CLEM), we capture and resolve early intermediates of starvation-induced ERphagy. These structures appear asymmetrically organized in representative images and are enriched in endogenous TEX264 and the ER-resident cargo reporter calnexin (CNX), together with the autophagy factors LC3 (Kabeya *et al*, 2000), p62 (Itakura & Mizushima, 2011), ubiquitin (Gonzalez *et al*., 2023). Their formation is substantially inhibited by pharmacologic inactivation of ULK1 kinase and of LC3 lipidation and in cells lacking the select autophagy components ATG9A, ATG13 or FIP200. Our findings provide an experimental strategy to enrich and morphologically study a normally transient, pre-lysosomal stage of ERphagy. They also reveal the architecture of the TEX264-positive ER membrane remodelling sites, where this process takes place and point to a restricted number of ER sites, at which ERphagy initiates.

## Results

### Generation and characterization of eTEX264-Halo MEF

To monitor the events of ERphagy induced by nutrient restriction that precede lysosomal delivery, we generated MEFs in which endogenous TEX264 was C-terminally fused to HaloTag (eTEX264-Halo; see Methods). Cells were labelled for 6h with 100nm Tetramethylrhodamine (TMR), a cell-permeable, fluorescent HaloTag ligand, and analyzed by CLSM. The TMR signal co-localized with the TEX264 immunoreactivity (**Fig. 1A**) and with the endogenous ER and nuclear envelope marker CNX (**Fig. 1B**), confirming that eTEX264-Halo retains the subcellular distribution reported for endogenous TEX264 (An *et al*., 2019; Chino *et al*., 2019).

**Fig. 1.**
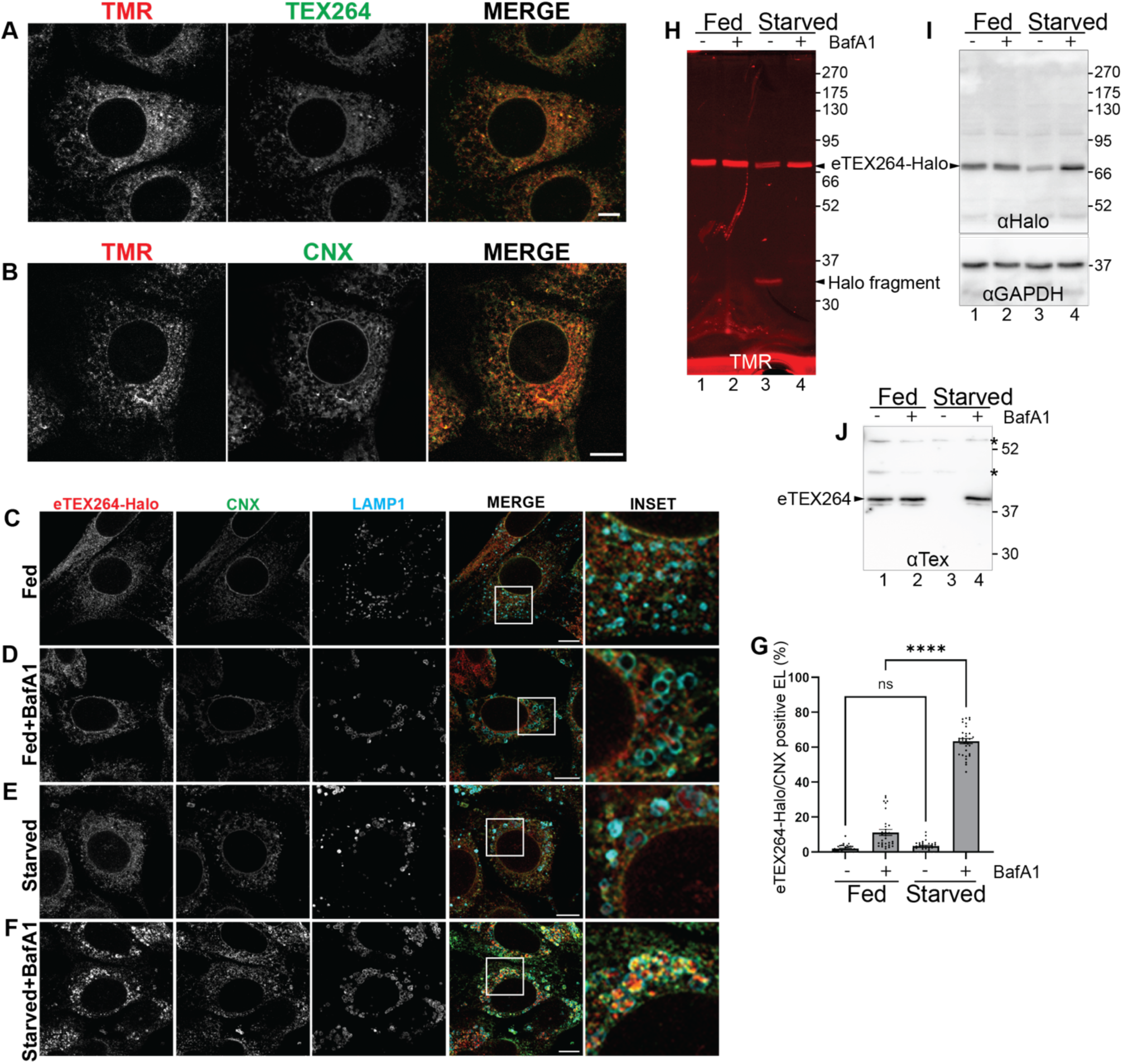
Endogenous (e)TEX264-Halo reports on starvation-induced ERphagy. **A** CLSM showing co-localization of the TMR signal with endogenous TEX264 and **B** co-localization of the TMR signal with the endogenous ER protein CNX. **C-D** CLSM analyses of endogenous CNX and eTEX264-Halo delivery to LAMP1 positive endolysosomes in fed conditions, ± 100nM BafA1. **E-F** same as **C-D** in starved cells. **G** Quantification of (**C-F**) by LysoQuant (Morone *et al*, 2020), n=31, 32, 33, 33, for fed, fed+BafA1, starved and starved+BafA1, respectively. N=3 independent experiments. Mean ± SEM; unpaired, two-tailed t-test, ^ns^P>0.05, ****P<0.0001. **H** Lysates of eTEX264-Halo MEF incubated with 100nM TMR in fed (lanes 1-2) or starved conditions (lanes 3-4) and 100nM BafA1 (lanes 2 and 4). The TMR fluorescence is seen in the gel. **I** The same gel has been transferred on PVDF membrane, which has been probed with αHalo to detect eTEX264-Halo (top panel) and αGAPDH antibodies (loading control, lower panel). **J** Immunoblot of MEF lysates with αTEX264 antibodies. *Cross reacting polypeptide bands. Scale bars 10µm.

Under nutrient rich conditions, only negligible quantity of the eTEX264-Halo and of CNX were detected within LAMP1-positive endolysosomes either in the absence or presence of Bafilomycin A1 (BafA1), which inhibits lysosomal acidification and thereby prevents the clearance of the cargo delivered within the degradative compartments (Klionsky *et al*, 2008) (**Figs. 1C, 1D**, respectively, quantification in **1G**). This is consistent with the low basal ERphagy activity in MEFs.

Nutrient restriction strongly stimulates ERphagy, with TEX264 acting as the major ERphagy receptor in this response (An *et al*., 2019; Chino *et al*., 2019). Accordingly, starvation in Earle’s Balanced Salt Solution (EBSS), which lacks amino acids and serum, markedly increased the accumulation of both eTEX264-Halo and CNX within LAMP1-positive endolysosomes in cells exposed to BafA1 (**Figs. 1E** *vs.* **1F**, **1G**), demonstrating enhanced delivery of ER material to degradative compartments.

We independently monitored ERphagy using a biochemical HaloTag processing assay, which exploits the generation of a protease-resistant HaloTag fragment following lysosomal degradation of Halo-tagged proteins (Rudinskiy *et al*., 2022; Rudinskiy *et al*., 2024; Yim *et al*., 2022). eTEX264-Halo MEF were labelled with TMR and cultured for 6h under either fed (**Figs. 1H**, **1I**, lanes 1 and 2) or starved conditions (lanes 3 and 4), in the absence (lanes 1, 3) or presence (lanes 2 and 4) of BafA1. Full-length eTEX264-Halo migrated just above 66kDa molecular weight marker (**Fig. 1H**, lanes 1-4), whereas a fluorescent HaloTag fragment of approximately 34kDa was detected specifically in starved cells (**Fig. 1H**, lane 3). This fragment corresponds to free HaloTag remaining after lysosomal degradation of the TEX264-containing ER material and therefore provides a biochemical read out of ERphagy flux (Rudinskiy *et al*., 2022; Rudinskiy *et al*., 2024; Yim *et al*., 2022). Consistent with its lysosomal origin, BafA1 abolished the generation of the free HaloTag fragment and stabilized full-length eTEX264-Halo (**Fig. 1H**, **1I**, lane 4). Together, these imaging and biochemical analyses establish that eTEX264-Halo faithfully recapitulates the localization (An *et al*., 2019; Chino *et al*., 2019; Kucinska *et al*., 2023), and lysosomal turnover of endogenous TEX264 (arrow eTEX264 in **Fig. 1J**), whose removal is induced upon starvation-induced ERphagy (lane 1 *vs.* 3) and is efficiently inhibited upon cell exposure to BafA1 (lane 4).

### Low temperature uncouples starvation-induced focal accumulation of eTEX264-Halo from lysosomal delivery

Nutrient restriction induces TEX264-positive puncta (clusters) that associate with autophagy factors and LC3 before TEX264-containing ER material reaches LAMP1-positive degradative compartments (An *et al*., 2019; Chino *et al*., 2019). Receptor clustering is emerging as a common principle for mitophagy (Wei *et al*, 2024) and ERphagy receptors activation (Foronda *et al*., 2023; Gonzalez *et al*., 2023) that promotes recruitment of the autophagy machinery at organelle membranes. For example, analyses of the ERphagy receptor FAM134B showed that its local clustering guided by starvation-induced post-translational modifications defines sites of ER remodelling, engagement of the autophagy machinery, and autophagosome initiation (Foronda *et al*., 2023; Gonzalez *et al*., 2023). Because these early steps of autophagy are transient under physiological conditions, we sought an experimental approach that would arrest ERphagy before lysosomal delivery and thereby enable the characterization of its early intermediates. Autophagic sequestration is strongly temperature-dependent and largely absent below 20°C in hepatocytes (Gordon *et al*, 1987). We tested whether lowering the temperature could temporally and spatially resolve starvation-induced ERphagy. Starvation experiments performed at 25°C showed a substantial inhibition of autophagic cargo delivery within LAMP1-positive degradative endolysosomes. At 25°C, in nutrient rich medium eTEX264-Halo and CNX retained their characteristic distribution throughout the ER and the nuclear envelope (**Figs. 2A-2B**) but nutrient deprivation induced the redistribution of eTEX264-Halo and CNX into cytosolic puncta approximately 250-1000nm in diameter (white arrowheads in **Figs. 2C-2D**). The formation of these eTEX264-Halo^+^/CNX^+^ puncta was unaffected by BafA1 indicating that their accumulation did not result from inhibition of lysosomal degradation (**Figs. 2C**, **2D**). Similar results were obtained in a cell line expressing the endogenously tagged ERphagy receptor eSEC62-Halo (Material and Methods and **Suppl. Fig. 1**).

**Fig. 2.**
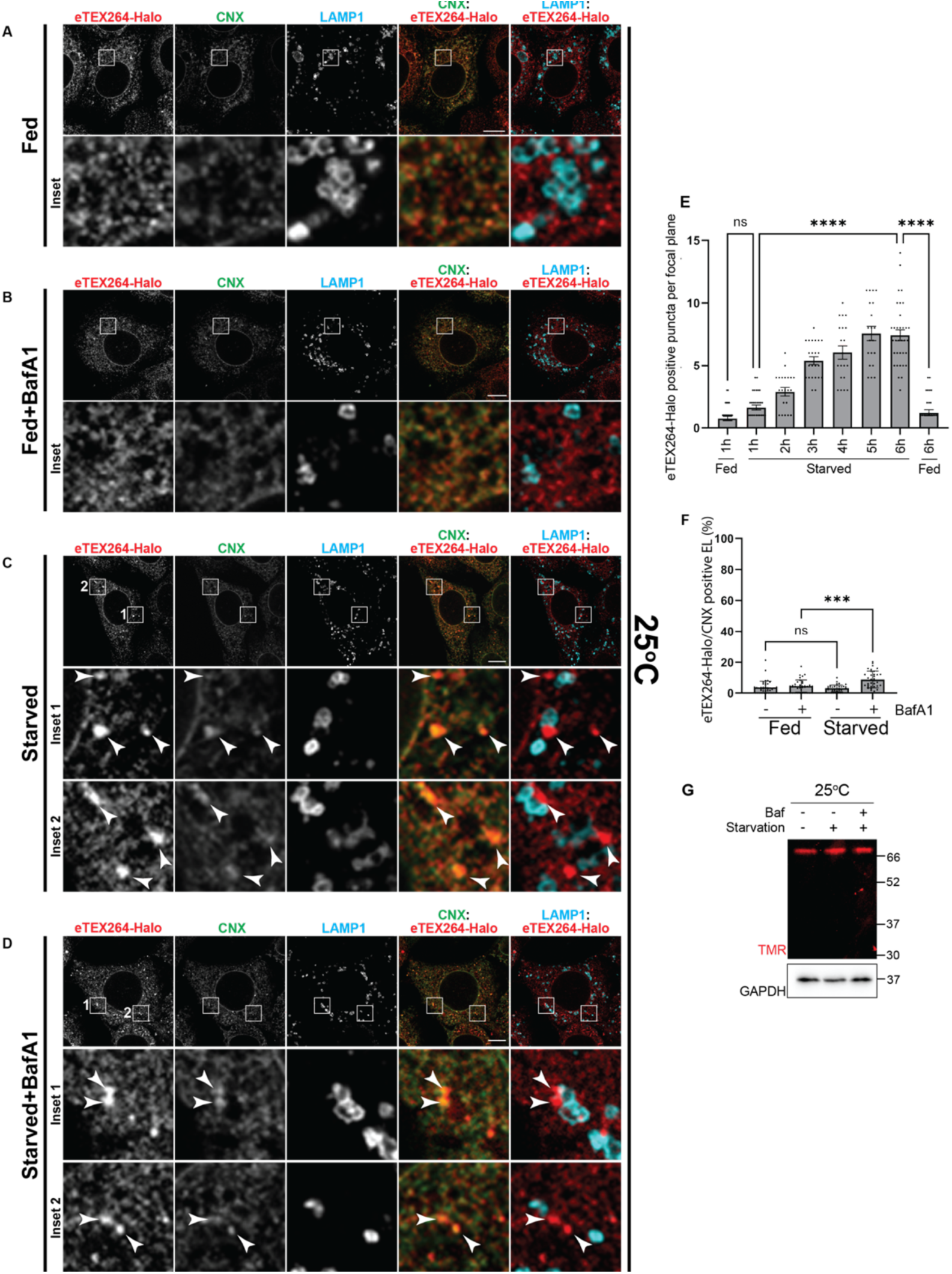
Low temperature uncouples starvation-induced focal accumulation of eTEX264-Halo and lysosomal delivery. **A-D** CLSM analysis of intracellular distribution of eTEX264-Halo at 25°C (fed or starved conditions, ± BafA1 100nM). The cytosolic eTEX264-Halo puncta formed in starved cells are marked with arrowheads. **E** Quantification of eTEX264-Halo positive puncta per cell/confocal plane starting from t=1h until t=6h at 25°C, in fed and in starved cells , n= 31, 30 for fed 1h, 6h, respectively. n= 31, 22, 23, 20, 20, 34 for 1h, 2h, 3h, 4h, 5h and 6h starved, respectively. N=3 independent experiments. A one-way ANOVA with Dunnett’s multiple comparisons test, ^ns^P>0.05; ****P<0.0001. **F** Quantification of eTEX264-Halo (**A-D**) by LysoQuant. For eTEX264-Halo/CNX-positive EL, n=40, 37, 35 and 35 for fed, fed+BafA1 100nM, starved and starved+BafA1 100nM respectively. N=3 independent experiments. Mean ± SEM; unpaired, two-tailed t-test, ^ns^P>0.05, ***P<0.001 **G** Lysates of 100nM TMR-labelled endogenous tagged eTEX264-Halo cells cultured with HEPES-buffered DMEM 10%FBS, EBSS or EBSS plus BafA1 100nM, at 25°C. Scale bar: 10µm

The number of eTEX264-Halo puncta progressively increased during the first 5h of starvation to reach a plateau of approximately 7 puncta per confocal plane with no further increase upon prolonged starvation (**Fig. 2E**, and **Suppl. Fig. 1** for eSEC62-Halo). At 25°C, the exposure of cultured cells to nutrient restriction induced the clustering of ERphagy receptors and of ERphagy cargo consistent with induction of ERphagy, but their subsequent lysosomal delivery within LAMP1-positive endolysosomes that occurs at 37°C (**Figs. 1F, 1G**) was strongly impaired (**Figs. 2C-2D**, **Fig. 2F**), consistent with a temperature block in the ERphagy program. Biochemical analysis corroborated the imaging data. Starvation at 25°C failed to generate the free HaloTag fragment characteristic of the lysosomal degradation of eTEX264-Halo (**Fig. 2G**, lane 2, and **Suppl. Fig. 1** for eSEC62-Halo), in contrast to the robust HaloTag processing observed during starvation-induced ERphagy at 37°C (**Fig. 1H**, lane 3). Thus, nutrient restriction at 25°C triggers the formation of TEX264- and CNX-containing puncta but the low temperature prevents their efficient delivery to lysosomes.

### eTEX264-Halo puncta formed at 25°C are competent for lysosomal delivery upon rewarming

We next asked whether the population of eTEX264-Halo puncta accumulated at 25°C can still reach LAMP1-positive endolysosomes once the permissive temperature is restored. Cells maintained in fed medium showed no induction of eTEX264-Halo or CNX puncta at 25°C (**Fig. 3A**) and negligible delivery of ER portions to LAMP1-positive endolysosomes after shift to 37°C (**Figs. 3C, 3E, 3H**). Formation of the eTEX264-Halo^+^/CNX^+^ puncta at 25°C required cell exposure to nutrient restriction (**Fig. 3B**, arrowheads). Under these conditions, eTEX264-Halo^+^/CNX^+^ puncta were mostly seen outside the LAMP1-positive endolysosomes (**Figs. 3B**, arrowheads, **3H**). Returning these cells to 37°C caused a rapid loss of cytosolic eTEX264-Halo^+^/CNX^+^ puncta (**Fig. 3G**) and their concomitant appearance within LAMP1-positive endolysosomes, where they accumulated in the presence of BafA1 (**Figs. 3D, 3F**, arrowheads, **3H**).

**Fig. 3.**
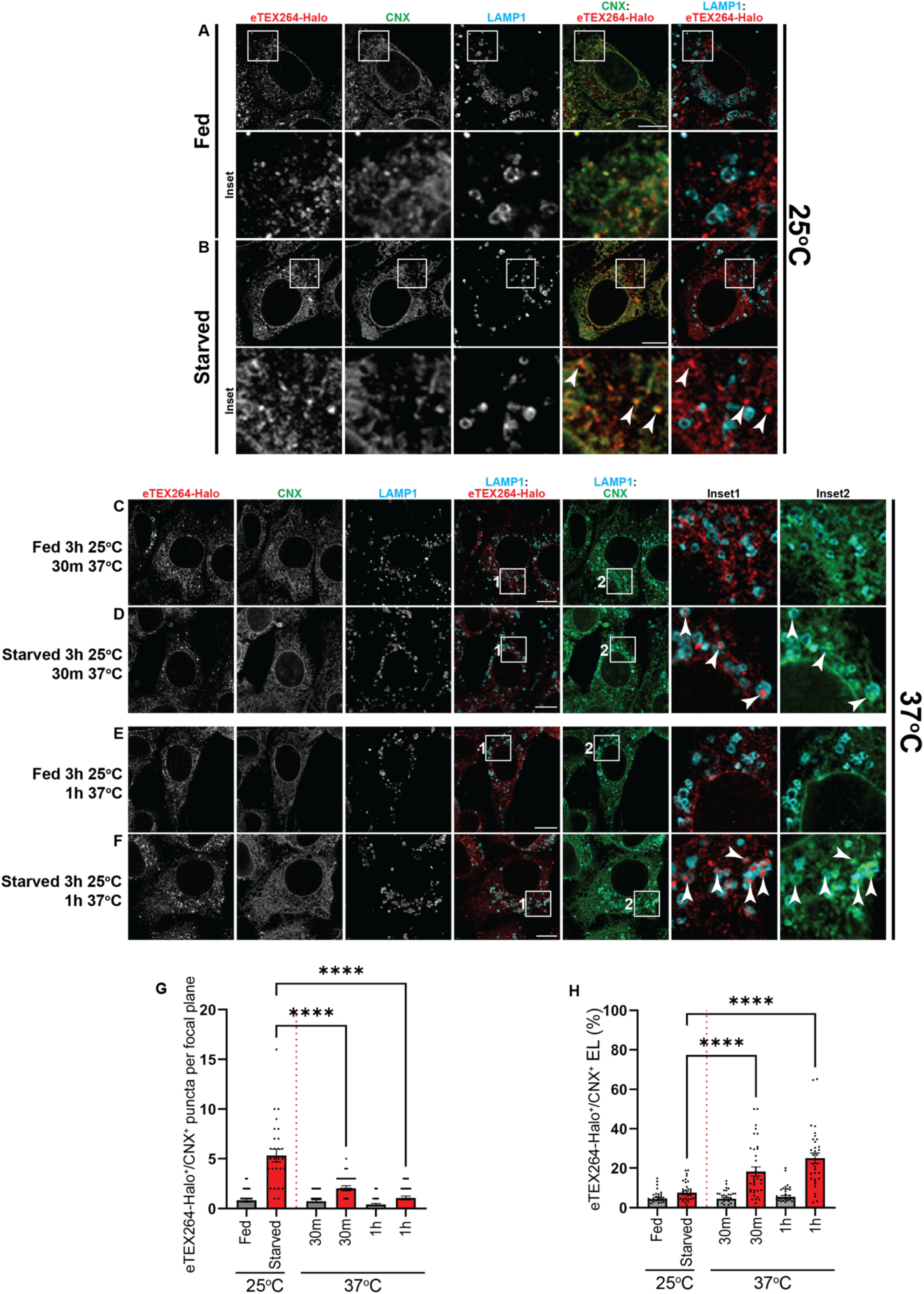
eTEX264-Halo puncta are delivered to endolysosomal compartments, after temperature switch. **A-B** CLSM analysis of TMR-labelled eTEX264-Halo cells kept in fed or starved conditions at 25°C for 3h (BafA1 100nM). **C, E** TMR-labelled eTEX264-Halo cells kept in fed conditions at 25°C for 3h were subsequently incubated for 30 (panel **C**) or for 60 min (panel **E**) at 37°C (BafA1 100nM). **D, F** Same as **C, E**, for starved cells. **G** Quantification of eTEX264-Halo^+^/CNX^+^ puncta/confocal plane in **A-F**. n=34, 34, 34 for fed cells and n= 28, 31, 31 for starved cells, respectively. N=3 independent experiments. Mean± SEM; A one-way ANOVA with Dunnett’s multiple comparisons test, ****P<0.0001. **H** Quantification with LysoQuant of (**A-F**). n=35, 34, 37 for fed cells and 35, 38, 33 for starved cells, respectively. Mean± SEM; N=3 independent experiments. A one-way ANOVA with Dunnett’s multiple comparisons test, ^ns^P>0.05, ****P<0.0001. Scale bar: 10µm

Live-cell imaging further indicated that delivery can resume upon rewarming of the cell culture media. In a recording in which the temperature of the culture medium was monitored throughout, individual eTEX264-Halo puncta entered LAMP1-GFP^+^ endolysosomes as the medium warmed from the 25 to about 27°C (arrowheads **Fig. 4**, **Movie 1**). Together, these data show that nutrient restriction at 25°C drives the focal accumulation of eTEX264-Halo puncta (**Figs. 2C, 2E, 3B**), whose delivery to endolysosomes is strongly reduced (**Figs. 2D, 2F, 2G**), and that this material is delivered within LAMP1-positive compartments once the temperature is raised (**Figs. 3D, 3F, 3H, 4**, **Movie 1**).

**Fig. 4.**
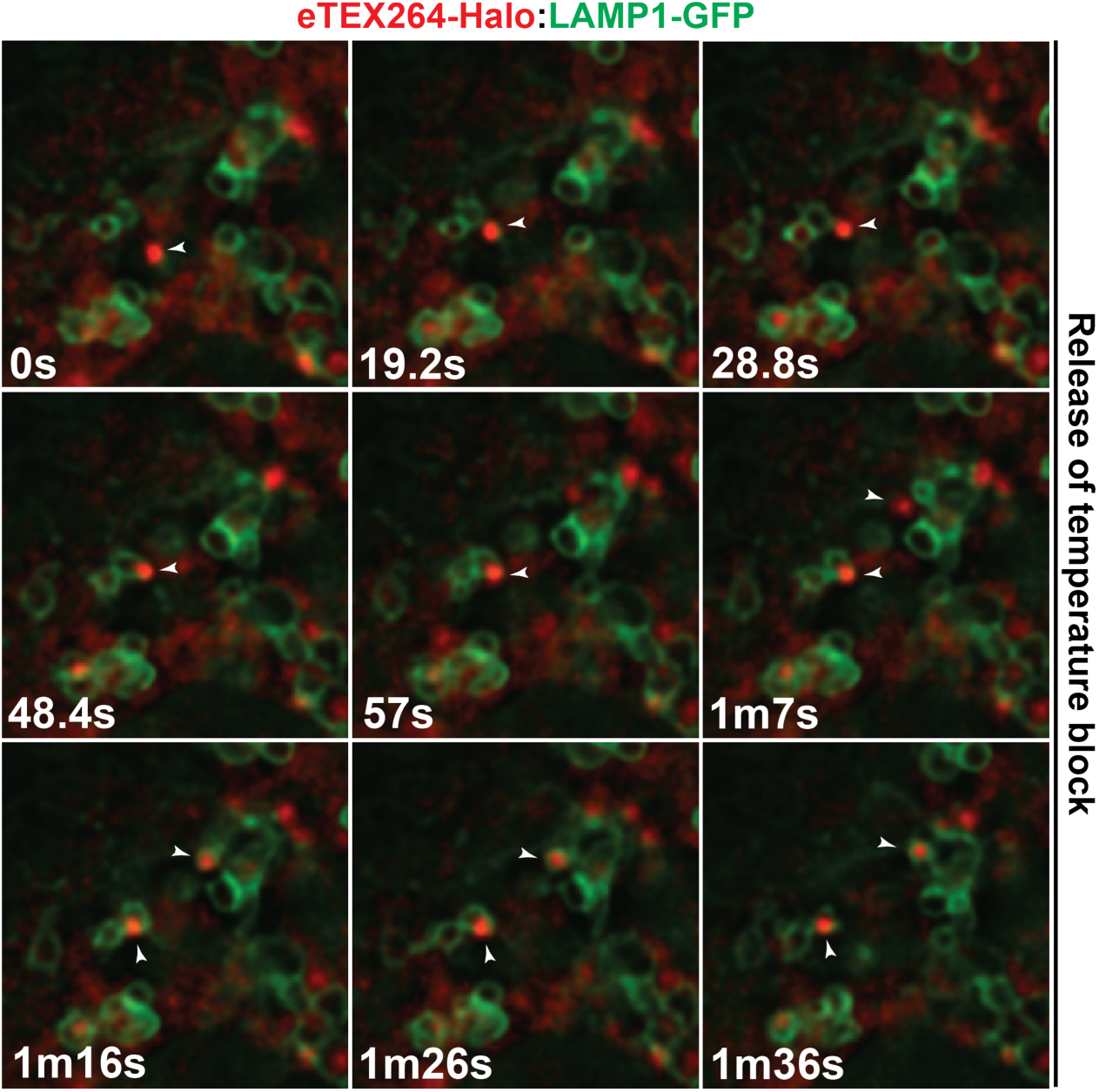
eTEX264-Halo puncta delivery within endolysosomal compartments is restored at permissive temperature. Representative frames of **Movie 1** showing delivery of individual eTEX264-Halo puncta (arrowheads) to LAMP1-GFP-positive endolysosomes after release of the 25°C temperature block (resumption of delivery block in live cell imaging checked in 2 independent experiments).

### eTEX264-Halo-positive ERphagy intermediates co-localize with autophagy markers

The reversible accumulation of eTEX264-Halo puncta at 25°C upon nutrient restriction provided an opportunity to characterize their ultrastructure. CLSM images show that eTEX264-Halo puncta formed at 25°C consistently co-localized with LC3 and p62/SQSTM1, as well as ubiquitin (**Fig. 5A**). At 37°C, where delivery to lysosomes is restored this autophagic cargo was delivered to LAMP1-positive compartments (**Fig. 5B**). The presence of cytosolic autophagic cargo indicates that starvation-induced eTEX264-Halo puncta accumulating at 25°C retain key molecular features of sites undergoing autophagosome capture despite their inability to complete lysosomal delivery.

**Fig. 5.**
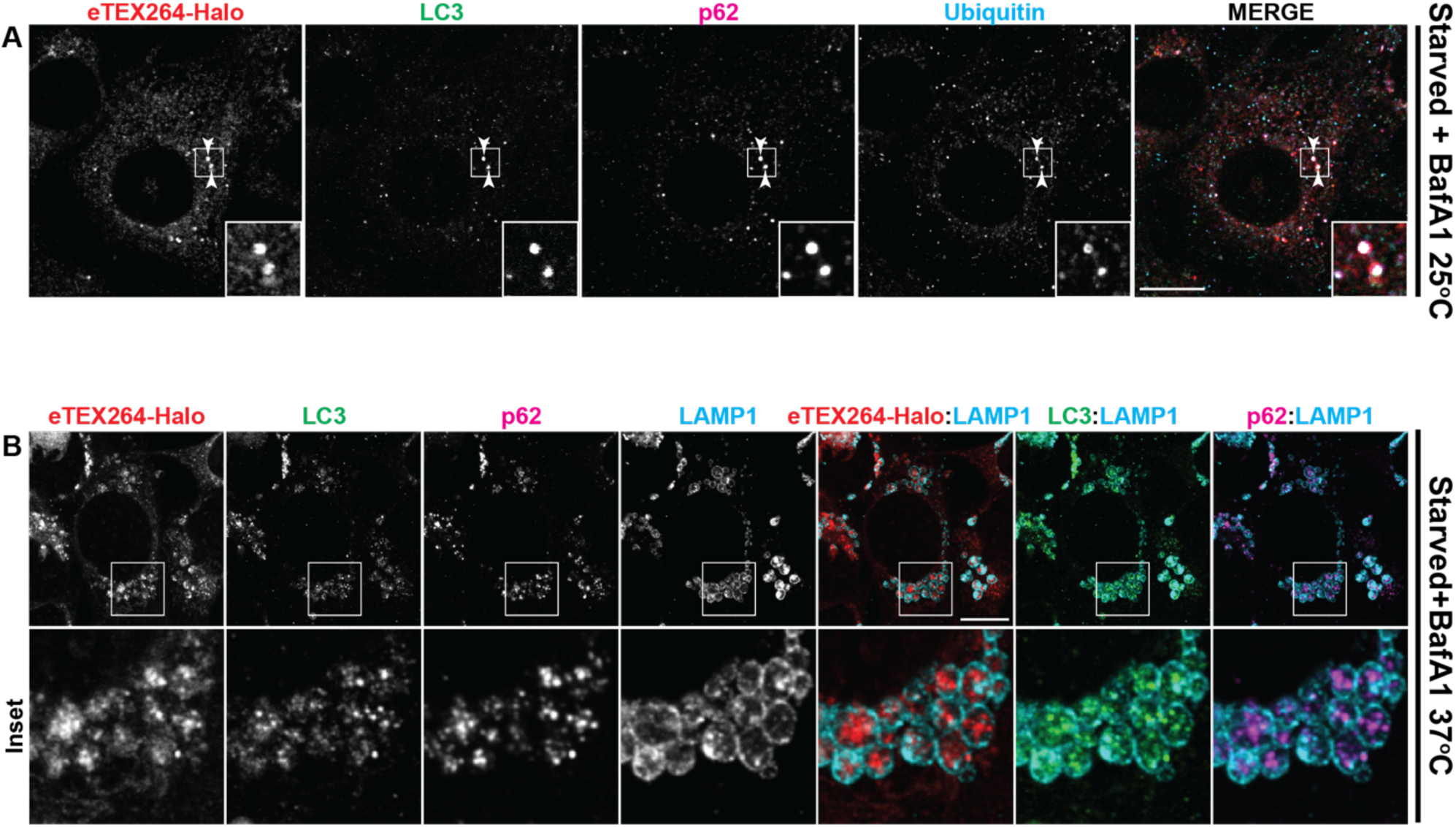
eTEX264-Halo puncta co-localize with autophagy cargo. (**A**) Representative images of CLSM analysis of eTEX264-Halo MEFs starved for 6h at 25°C in the presence of BafA1 and stained for LC3, p62, ubiquitin (representative of at least 3 independent experiments), (**B**) CLSM analysis of eTEX264-Halo MEFs starved for 6h at 37°C in the presence of 100nM BafA1. Stains are for LC3, p62 and LAMP1 (representative of at least 2 independent experiments). Scale bar: 10µm

### Formation of the temperature-arrested eTEX264-Halo puncta requires the core autophagy machinery

To assess the possibility that starvation-induced formation of eTEX264-Halo puncta requires the autophagosome biogenesis machinery, we first compared their formation upon cell starvation at 25°C (**Figs. 6A** *vs.* **6B**, **6G**), following pharmacological inhibition of ULK1/2 with MRT68921 that substantially hampers phagophore formation (**Figs. 6C** *vs.* **6D**, **6G**) (Petherick *et al*, 2015) and upon cell exposure to ATG7-IN-3, an ATG7 inhibitor (**Figs. 6E** *vs.* **6F**, **6G**) (Huang *et al*, 2020).

**Fig. 6.**
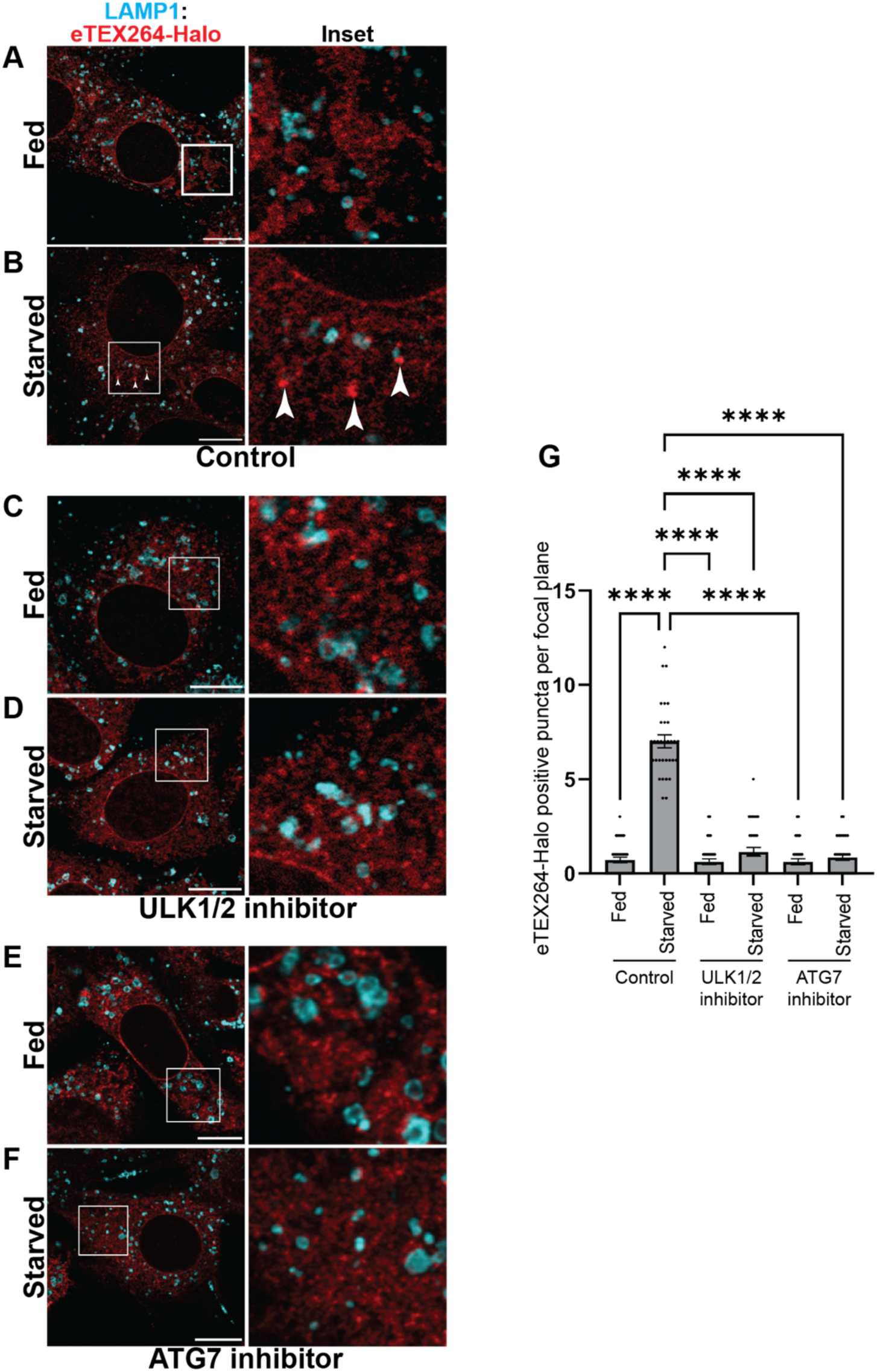
The core autophagosome biogenesis machinery is required for starvation-induced formation of eTEX264-Halo puncta at 25°C. **A-B** CLSM analysis of TMR-labelled eTEX264-Halo cells maintained under fed or starved conditions at 25°C for 6h. **C-D** Same as **A-B** in the presence of 1µM MRT68921. **E-F** Same as **A-B** in the presence of 10µM ATG7-IN-3. **G** Quantification of eTEX264-Halo puncta in **A-F**. n=35, 32, 34, 35, 34, 34, respectively. N=3 independent experiments. Mean± SEM; one-way ANOVA with Dunnett’s multiple comparisons test, ****P<0.0001.

The treatment of eTEX264-Halo MEFs with MRT68921 substantially reduced the formation of starvation-induced eTEX264-Halo puncta at 25°C (**Figs. 6B** *vs.* **6D, 6G**) supporting the involvement of the ULK1/ULK2 complex in the focal accumulation of the ERphagy receptor in response to an exogenous signal that activates ERphagy. Likewise, cell exposure to ATG7-IN-3 substantially reduced formation of eTEX264-Halo puncta (**Figs. 6B** *vs.* **6F, 6G**), supporting a requirement for the ATG7-dependent conjugation machinery in the focal accumulation of the ERphagy receptor under the temperature block.

To independently, and genetically test the requirement for the autophagosome biogenesis machinery, we analyzed U2OS cells (an established human bone mesenchymal tumor cell line (Ponten & Saksela, 1967)) lacking ATG9A, ATG13 or FIP200 (Mauthe *et al*, 2021). Starvation at 25°C induced the formation of eTEX264-Halo puncta and of endogenous p62^+^ puncta in wild-type U2OS cells (**Suppl. Figs. 2A** *vs.* **2B**, **2F**). Starved U2OS cells lacking ATG9A, a lipid scramblase required for the expansion of the phagophore membrane (Karanasios *et al*, 2016; Lin *et al*, 2025; Olivas *et al*, 2023; Orsi *et al*, 2012; Young *et al*, 2006), showed a significant reduction of eTEX264-Halo puncta (**Suppl. Figs. 2B** *vs.* **2C**, **2F**). The same result was obtained for U2OS lacking ATG13 (**Suppl. Figs. 2D, 2F**), and FIP200 (**Suppl. Figs. 2E, 2F**), two key components of the ULK complex which is responsible for initiating phagophore formation (Ganley *et al*, 2009; Hama *et al*, 2025; Hara *et al*, 2008; Jung *et al*, 2009). Together, these pharmacological and genetic results support a requirement for the autophagosome biogenesis and the ATG7-dependent conjugation machinery in the accumulation of starvation-induced eTEX264-Halo puncta at 25°C.

### Expansion microscopy reveals the organization of eTEX264-Halo-positive ERphagy intermediates

The requirement of an activating cue (nutrient restriction, **Figs. 2, 3**), the involvement of the autophagosome biogenesis machinery in their formation (**Fig. 6, Suppl. Fig. 2**), and the resumption of lysosomal delivery upon rewarming of the cell culture (**Figs. 3, 4**), suggested that eTEX264-Halo/CNX puncta represent specialized ER subdomains destined for lysosomal clearance during starvation-induced ERphagy. Because the 25°C temperature-block stabilizes these otherwise transient structures, we further examined their ultrastructure by expansion microscopy (ExM) combined with immunostaining. Our protocol expanded samples by 3.8 ± 0.2 fold in linear dimension, as estimated by comparing nuclear dimensions before and after expansion, in agreement with previous reports (Chen *et al*, 2015; Gambarotto *et al*, 2019; Louvel *et al*, 2023).

In starved cells at 25°C, these putative ERphagy initiation sites consisted of curved, eTEX264-Halo-positive ER membranes that, consistent with CLSM images (**Figs. 2-5**), also contained CNX (**Fig. 7A**, arrows), LC3 (**Fig. 7B**, arrows) and were partially positive for ubiquitin (**Fig. 7C**, red arrows). The expansion of the puncta revealed that p62 (seen as a co-localized protein in CLSM, **Fig. 5A**) was also abundant at these sites but did not overlap with the ER markers. Instead, it occupied a discrete domain directly adjacent to the eTEX264-Halo/CNX-positive membranes (**Fig. 7A**, blue arrows). We refer to this polarized arrangement, with ER membrane on one side and a p62-positive domain on the other, as asymmetric.

**Fig. 7.**
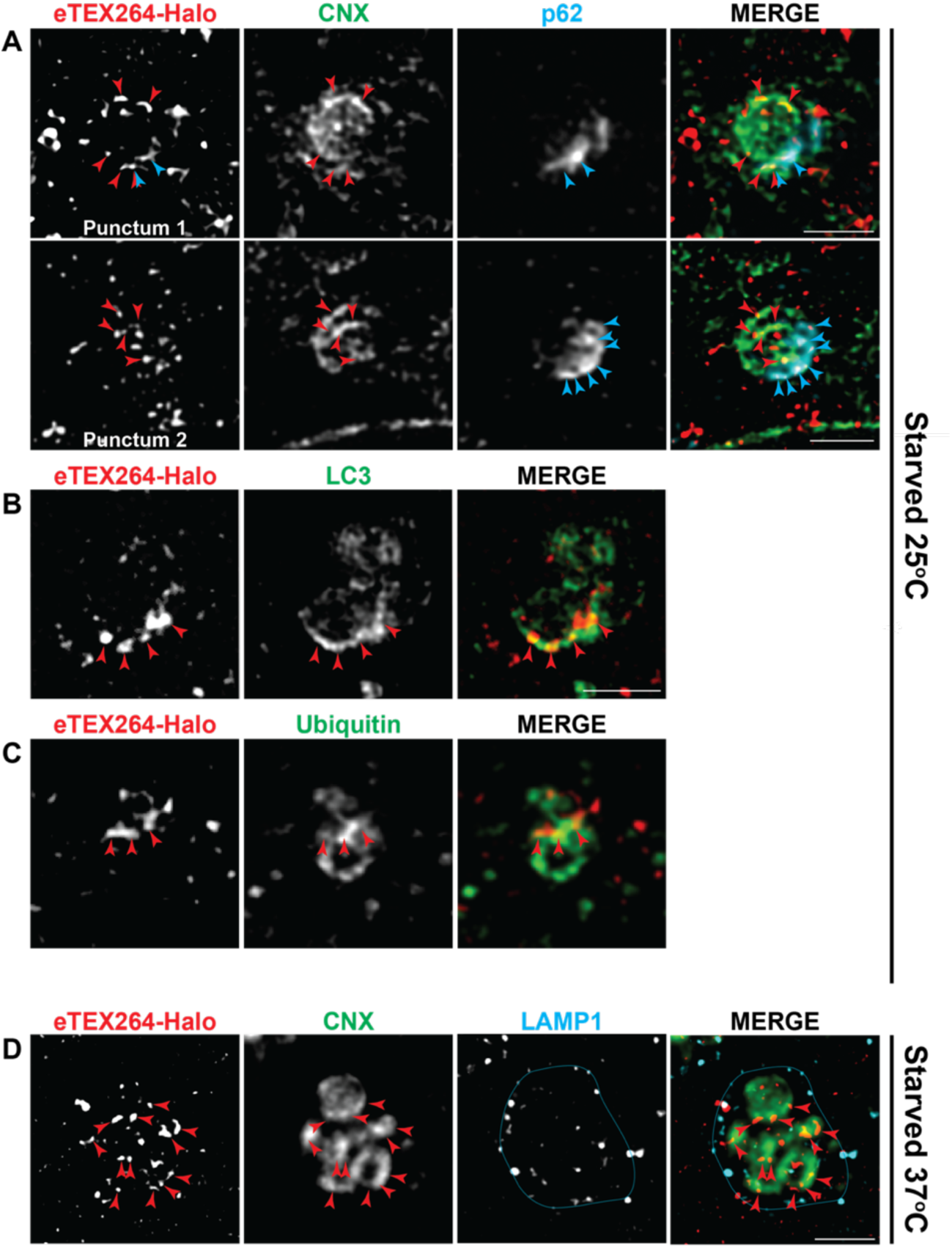
Ultrastructure of starvation-induced eTEX264-Halo puncta in ExM. **A** ExM micrographs of eTEX264-Halo puncta at 25°C showing CNX-positive ER membrane on one side and p62-positive material on the other side of two distinct puncta selected as examples from 2 independent experiments. **B, C** Same as **A** for LC3 and ubiquitin, respectively. **D** ExM micrographs of the eTEX264-Halo puncta after delivery within LAMP1-positive endolysosomes at the permissive temperature of 37°C in the presence of BafA1. Scale bar: 2 µm. Scale bars refer to expanded sizes. These ExM images are representative for the puncta ultrastructure of 3 biological replicates for the co-localization shown in panel **A**, of 2 biological replicates for panels **B** and **C** (which confirm the CLSM analysis in Fig. 6A). The representative LAMP1-positive endolysosome containing ER material is the ExM image of cells from 1 experiment (delivery of ER portions within LAMP1-positive endolysosomes at 37°C is shown in Figs. 1F, 1G, 3F, 3H**, 4, 6E** and **Movie 1**).

As shown above, the ExM images confirmed that in cells starved at 37°C in the presence of BafA1, CNX- and eTEX264-Halo-positive ER material is delivered to the LAMP1-positive compartments (**Fig. 7D**, red arrows, LAMP1 boundary, blue line). This is consistent with completed lysosomal delivery under permissive conditions.

### Electron tomography reveals ER membrane remodelling at eTEX264-Halo-positive sites

To define the membrane architecture underlying the starvation-induced eTEX264-Halo-positive ERphagy intermediates, we analyzed their structures by correlative light and electron microscopy (CLEM). eTEX264-Halo MEFs were starved at 25°C, eTEX264-Halo puncta were located by fluorescence microscopy, and the corresponding regions were then analyzed by electron tomography (**Fig. 8A)**. Tomograms (**Fig. 8B**) and the derived 3-D reconstructions (**Fig. 8C**) showed gold-labelled eTEX264-Halo-positive ER membranes organized into curved, cradle-like structures consistent with the morphology observed by ExM (**Fig. 7**).

**Fig. 8.**
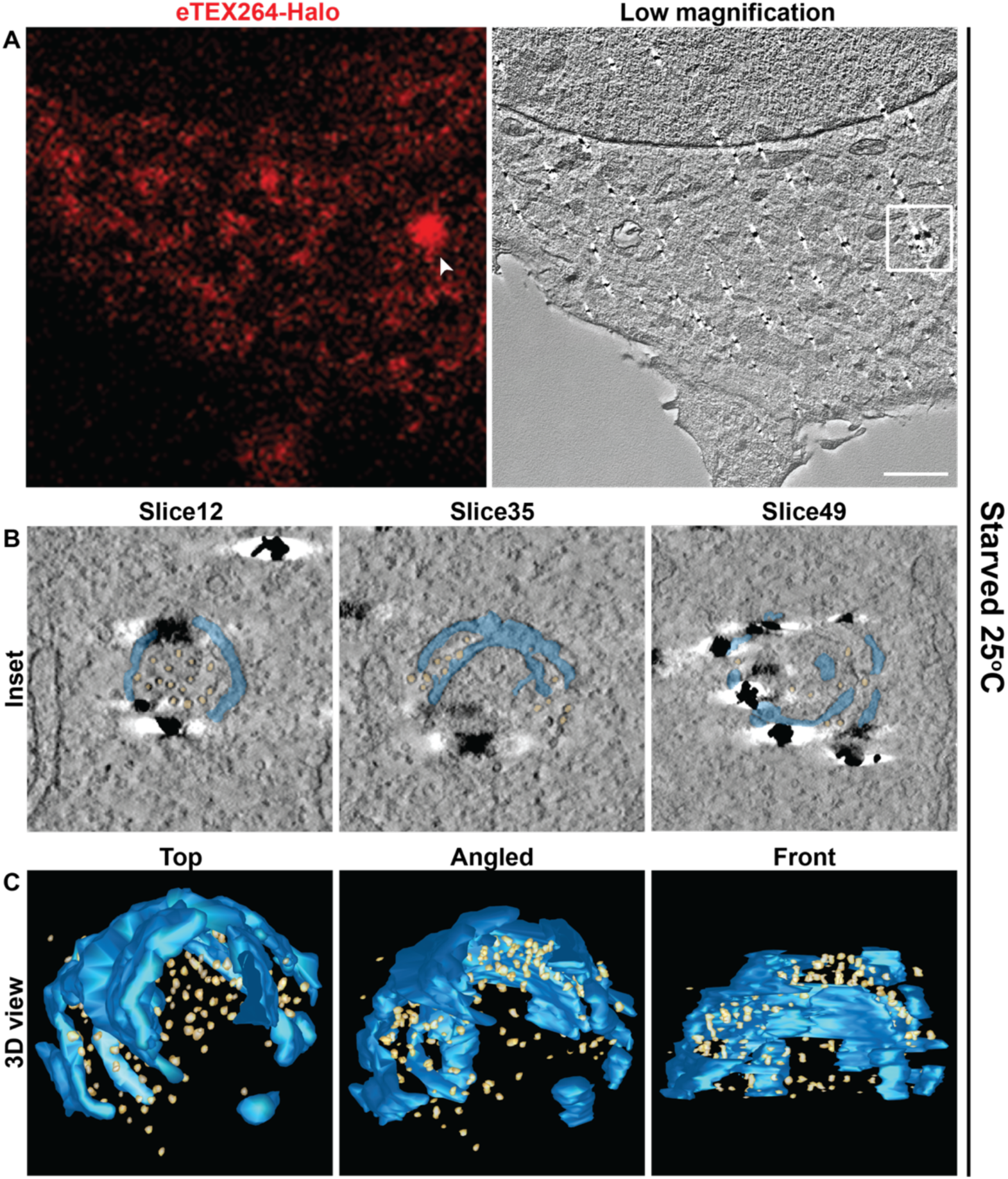
Electron tomography reveals membrane remodelling at eTEX264-Halo puncta. **A** CLEM analysis of TMR-labelled eTEX264-Halo cells starved at 25°C for 6h showing the chosen CLSM field and the corresponding EM field **B** Selected slices of electron tomography images showing gold-labelled, eTEX264-Halo-positive ER membranes (blue), vesicles (yellow) **C** 3D reconstruction of (**B**). Scale bar and insets: 1µm.

Small vesicular profiles were present within and adjacent to the concave face of the cradle (**Fig. 8B**, slices 12, 35, and 49). In addition, a double-membrane structure protruded from eTEX264-Halo-positive ER membranes (highlighted in blue, **Fig. 8B**, slice 35), and further gold-positive ER portions were found close to the opening of the curved ER domain (**Fig. 8B**, slice 49). These features are in line with recent ultrastructural observations obtained in cells overexpressing the ERphagy receptor FAM134B that phagophore membranes sequestering ER subdomains emerge directly from remodeled ER (Ma *et al*, 2026).

This ultrastructural organization identifies eTEX264-Halo puncta as highly organized sites of ER membrane remodelling associated with nascent autophagic membranes that capture cytoplasmic material to be removed from cells. Arresting the pathway at 25°C therefore renders an otherwise transient ERphagy intermediate accessible to ultrastructural analyses and reveals that eTEX264-Halo-positive ER domains adopt a curved, cradle-like organization reminiscent of the ER subdomains described as sites of autophagosome biogenesis (Axe *et al*, 2008; Hayashi-Nishino *et al*, 2009).

## Discussion

ERphagy, the selective autophagic clearance of ER subdomains, proceeds through activation of ER-resident receptors that, upon post translational modification of their cytosolically exposed regions assemble into membrane-associated platforms. These platforms remodel the ER membrane, engage the core autophagy machinery, drive physical separation of the selected ER subdomain, and ultimately deliver this material to the degradative endolysosomes (Gubas & Dikic, 2022; Iavarone *et al*, 2022; Reggiori & Molinari, 2022). TEX264 is the principal receptor for starvation-induced ERphagy in mammalian cells, and its role in delivering ER material to degradative compartments is established (An *et al*., 2019; Chino *et al*., 2019). Autophagosome biogenesis events are completed within minutes and occur at only a handful of sites per cell (Koyama-Honda *et al*., 2013), making the early, receptor-driven steps that precede lysosomal delivery exceptionally difficult to resolve in unperturbed cells. Existing strategies to enrich early intermediates by genetically inactivate the autophagic program accumulate structures irreversibly and therefore cannot establish whether the trapped material remains competent for lysosomal delivery and clearance. Here, by combining endogenous HaloTag knock-in with a reversible 25°C temperature block, we bypass these limitations to monitor the time-dependent, starvation-induced, focal accumulation of TEX264. The rapid resumption of lysosomal delivery upon rewarming at 37°C, and above about 27°C in live imaging, demonstrates that eTEX264-Halo puncta are genuine, on-pathway intermediates of the ERphagy program. That receptor’s focal accumulation, cargo (CNX) segregation, ER membrane remodelling all proceed at 25°C, whereas delivery to endolysosomes is substantially inhibited, places the temperature-sensitive step downstream of phagophore nucleation rather than at the initial sensing of nutrient stress or the assembly of receptor platforms. The block is neither receptor- nor cell-type-specific because starvation at 25°C uncoupled ERphagy receptor clustering from lysosomal delivery of ER portions also in cells expressing the endogenously tagged version of the ERphagy receptor SEC62 (**Suppl. Fig. 1**) and induced puncta of endogenous, untagged TEX264 in U2OS cells (**Suppl. Fig. 2**). The reduction of eTEX264-Halo puncta upon ULK1/2 or ATG7 inhibition and in cells lacking ATG9A, ATG13 and FIP200 (**Fig. 5, Suppl. Fig. 2**) shows that their accumulation is coupled to the core autophagosome biogenesis machinery. The cradle-like organization revealed by ExM and electron tomography, curved eTEX264-Halo- and CNX-positive ER membranes that are near a discrete p62-containing domain at the concave opening suggests that ERphagy receptors may nucleate spatially compartmentalized platforms for phagophore formation at the ER surface, in which cargo segregation and omegasome-associated membrane remodelling occur at discrete but adjacent subdomains rather than within a single homogeneous assembly.

Two limitations of this study should be noted. First, we do not identify the molecular basis of the 25°C block. Our data place it downstream of phagophore nucleation but do not distinguish among phagophore closure, autophagosome maturation and fusion with endolysosomes, and we therefore refrain from assigning it to a specific step. The arrested structures are best understood as pre-delivery intermediates. Second, low temperature slows many cellular processes, and we have not excluded contributions from reduced membrane fluidity, altered motor activity or reduced ATP availability. Relevant precedents indicate that autophagosome-lysosome fusion is itself temperature-dependent in vitro (Koga *et al*, 2010) and that autophagic sequestration is strongly temperature-dependent and largely absent below 20°C in hepatocytes (Gordon *et al*., 1987).

This gap does not limit utility of the block. Temperature blocks have long served to dissect membrane trafficking (Griffiths *et al*, 1989; Griffiths *et al*, 1985; Kuismanen & Saraste, 1989; Matlin & Simons, 1983) precisely because they arrest specific, reproducible steps without requiring prior knowledge of their molecular basis. The 15°C and 20°C blocks that retain secretory cargo in the ER-Golgi intermediate compartment and in the trans-Golgi network, respectively, became standard tools (Matlin & Simons, 1983; Saraste & Kuismanen, 1984) even though their mechanistic basis remains incompletely understood (Gilbert *et al*, 2018; Martínez-Alonso *et al*, 2007). What validated them was reproducibility, specificity and reversibility. The 25°C block satisfies the same criteria as it operates in different cell lines, with endogenous and ectopically expressed protein reporters, and is rapidly reversed upon rewarming, with delivery resuming above 27°C (**Figs. 3, 4**). Defining its mechanistic basis, including possible off-target effects as documented for the classical secretory block (Gilbert *et al*., 2018; Martínez-Alonso *et al*., 2007), is a direction for future work. In the meantime, it provides a means of synchronizing autophagic pathways and of capturing intermediates of the programs that would otherwise be very difficult if not impossible to characterize both morphologically or mechanistically.

## Supplementary figures

**Suppl. Fig. 1.**
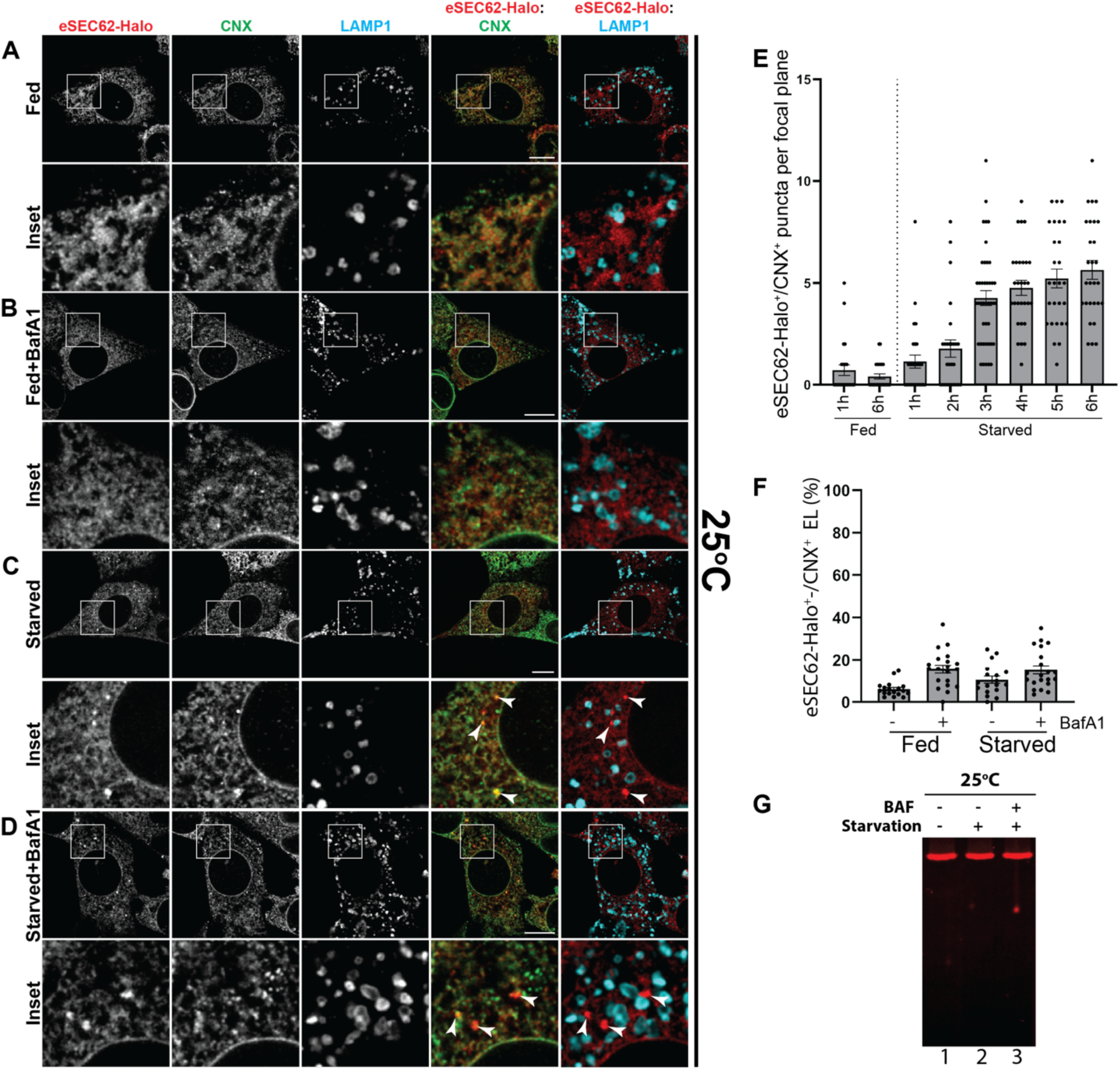
Low temperature uncouples eSEC62-Halo clustering and lysosomal delivery. **A-D** CLSM analysis of eSEC62-Halo and endogenous CNX delivery within LAMP1 positive endolysosomes (fed or starved conditions, ± BafA1 100nM at 25°C). eSEC62-Halo puncta are marked with arrowheads. **E** Quantification of eSEC62-Halo positive puncta per cell/confocal plane starting from t=1h until t=6h, without or with starvation (**A, C** panels respectively), n= 25 for 1h and n= 27 for 6h in fed conditions, n= 29 for 1h, n= 27 for 2h, n=42 for 3h, n= 29 for 4h, n= 27 for 5h, and n= 28 for 6h in starved conditions, N=2 independent experiments. A one-way ANOVA with Dunnett’s multiple comparisons test, nsP>0.05; ****P<0.0001. **F** Quantification of (**A-D**) by LysoQuant. For eSEC62-Halo/CNX positive EL, n=18, 20, 18 and 21 for fed, fed+BafA1 100nM, starved and starved+BafA1 100nM respectively. N=2 independent experiments. Mean ± SEM; unpaired, two-tailed t-test, ^ns^P>0.05 **G** Lysates of 100nM TMR-labelled eSEC62-Halo cells cultured with HEPES-buffered DMEM 10%FBS, EBSS or EBSS plus BafA1 100nM, at 25°C. Scale bar: 10µm.

**Suppl. Fig. 2.**
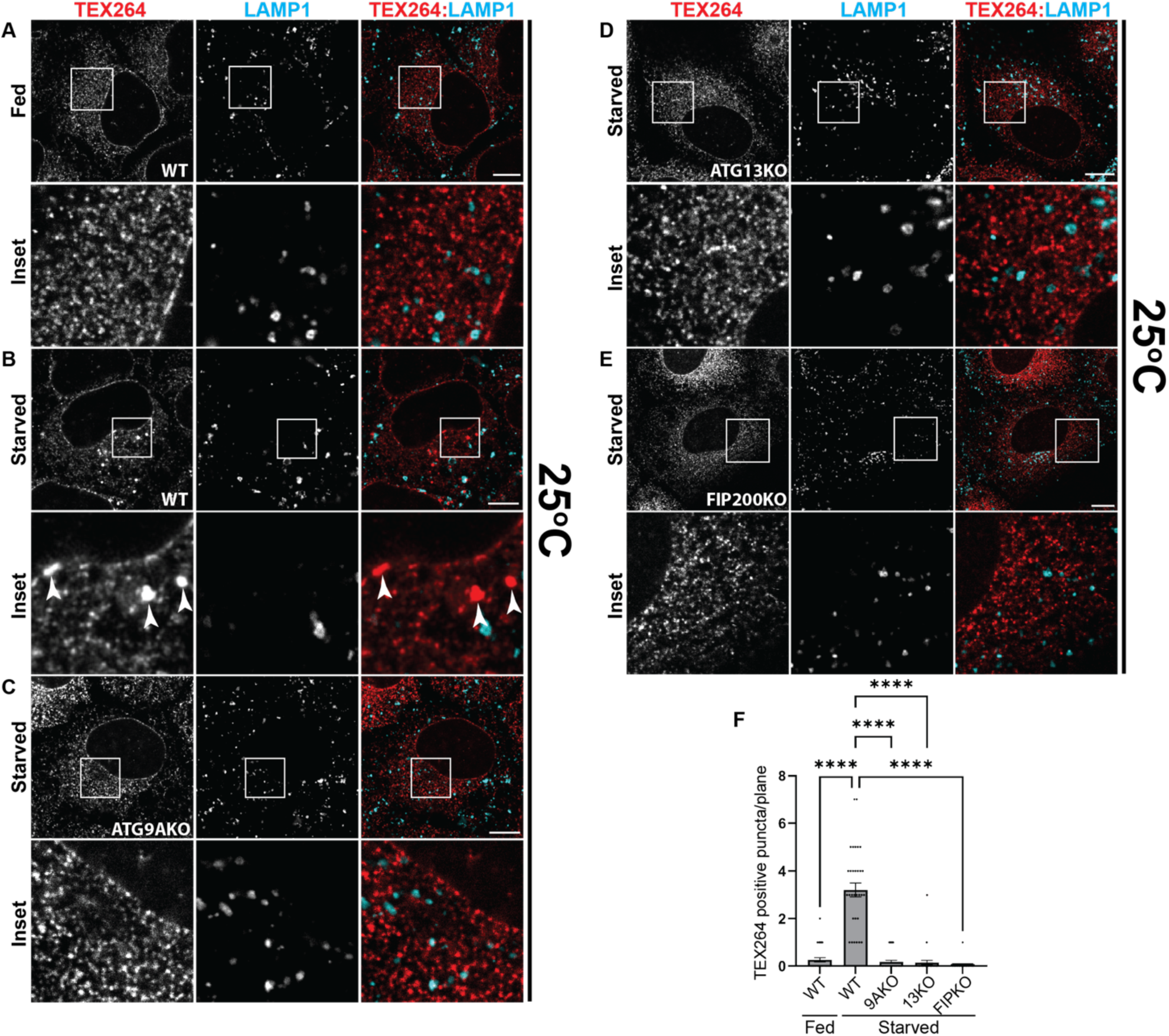
Starvation-induced endogenous TEX264 puncta at 25°C in WT, ATG9A-, ATG13- and FIP200-KO U2OS cells. **A** CLSM image of fed U2OS WT cells at 25°C **B** Same as **A** in starved U2OS WT cells. **C** Same as **B** in ATG9A-KO U2OS cells. **D** Same as **B** in ATG13-KO U2OS cells. **E** Same as **B** in FIP200-KO U2OS cells. **F** Quantification of eTEX264-Halo puncta per cell/confocal plane in (**A-E**), n=35, 34, 35, 35, 35 for U2OS WT fed and starved, starved ATG9A-KO U2OS, starved ATG13-KO U2OS, starved FIP200-KO U2OS, respectively. N=3 independent experiments. A one-way ANOVA with Dunnett’s multiple comparisons test, ****P<0.0001.

**Movie 1 eTEX264-Halo puncta delivery within endolysosomal compartments is restored at permissive temperature**

Movies were recorded on a Leica THUNDER microscope with a Leica HC PL APO 100x/1.44 OIL UV objective. Images were acquired every 9.6s, for a total length of the movie of 154s. The temperature of the culture medium was recorded throughout imaging using a semiconductor temperature probe immersed in the medium connected to a CENTER 522 Wireless Datalogger Hygro Thermometer, using the SE522 software (v 3.2.0.0).

## Material and Methods

### Generation of endogenous tagged eTEX264-Halo and SEC62-Halo cells and cell culture

TEX264-Halo MEF cells were generated by transfecting MEF WT cells with guideRNA-Cas9 plasmid pSpCas9(BB)-2A-Puro (PX459) V2.0 (Plasmid#62988 from Addgene) and a donor plasmid. gRNA targeting the C-terminal part of the protein was designed with the website https://crispr.mit.edu (5’-CACCGCATTACTCCTCACCCCTCTC-3’) and synthesized by Microsynth. The donor vector was designed by flanking the HaloTag sequence with two homology arms of 800 nucleotides each and cloned into a pUC19 plasmid (Genscript). Transfection was made using JetPrime (Polyplus) according to the manufacturer’s instructions. Two days after transfection, transfected cells were selected using 2µg/ml puromycin (Sigma), clonal populations were picked and cultured. Clones that integrated the HaloTag were selected based on their fluorescent signal using ImageXpress and screened by SDS-PAGE and Western blot. SEC62-Halo cells were generated following the same protocol of eTEX264-Halo MEF, using the oligonucleotide 5’-CACCGGAAATCATAATCTGACTAGC-3’ as gRNA. U2OS cells were a kind gift from Fulvio Reggiori. MEF cells were cultured in HEPES-buffered DMEM supplemented with 10% fetal calf serum (FCS, Gibco) at 37 °C. U2OS cells were cultured in HEPES-buffered DMEM supplemented with 10% fetal calf serum (FCS, Gibco) and 1mM sodium pyruvate at 37 °C in a 5% CO2 atmosphere.

### TMR labelling and cell lysis

Endogenous tagged eTEX264-Halo MEF cells were seeded on 6cm dishes. During treatment they were incubated with 100nM TMR HaloTag ligand (Promega) with or without 100 nM BafA1 (Invivogen) and then washed with ice-cold phosphate buffered saline (PBS) containing 20mM N-ethylmaleimide. Then they were lysed 20 min at 4°C in CHAPS buffer (2% CHAPS in HEPES-buffered saline (HBS), pH 6.8) or RIPA buffer (1%Triton X-100, 0.1% SDS, 0.5% sodium deoxycholate in HBS, pH 7.4), both with 20mM NEM and protease inhibitors. Dishes were scraped and the post-nuclear supernatants (PNS) were collected by centrifugation at 10000g for 10 min.

### Gel visualization and western blot

Samples were loaded on 10% SDS-PAGE (polyacrylamide gel electrophoresis) and run at 150V. Gels were first scanned for TMR fluorescence with the Amersham Typhoon scanner (Cytiva) using the 532nm laser and 570BP20 filter. Then gels were transferred onto polyvinylidene difluoride (PVDF) membranes with the Trans-Blot Turbo Transfer system (Bio-Rad). Membranes were blocked with 8% (w/v) non-fat dry milk (Bio-Rad) in Tris-buffered saline containing 1% Tween (TBS-T) and incubated with primary antibodies overnight, and for 45 min with Protein A HRP-conjugated or HRP-conjugated secondary antibodies. Membranes were developed using WesternBright Quantum (Advansta) and the signals were captured using Fusion FX imaging system (VILBER/Witec).

### Confocal laser scanning microscopy (CLSM)

Endogenous tagged eTEX264-Halo cells were seeded on alcian-blue coated coverslips. Cells were incubated for 45min at 25°C to allow the temperature to slowly decrease, cell medium was then exchanged to HEPES-buffered DMEM 10% FBS/EBSS for 6 hours with 100nM TMR HaloTag ligand.

Cells were treated with 10 µM ATG7-IN-3 or 1 µM MRT68921. Cells were then washed in PBS and fixed in 3.7% formaldehyde diluted in PBS at RT. Subsequently they were permeabilized in 0.05% saponin, 10% goat serum, 10mM HEPES and 15 mM glycine (PS) for 15 min. Cells were incubated with the primary antibodies diluted 1:100 in PS for 1h30min, then washed two times in PS and incubated with Alexa Fluor-conjugated secondary antibodies 1:300 in PS for 45min. Cells were washed three times in PS and mounted with Mount Liquid anti-fade (Abberior). Confocal images were acquired with Leica STELLARIS 5 and STELLARIS 8 with a Leica HCX PL APO lambda blue 63x/1.40 oil objective. The software used was Leica LAS X. Fluorescence emissions were collected at the following ranges: 430–490 nm (Alexa Fluor 405), 504–587 nm (Alexa Fluor 488), 592–644 nm (Alexa Fluor 568) and 658–750 nm (Alexa Fluor 647). Image analysis and quantification were performed using LysoQuant and ImageJ 2.16.0/1.54p software. Image post-processing was performed with Adobe Photoshop (v26.2.0).

### Live cell imaging

Endogenous tagged eTEX264-Halo cells were seeded on 35mm glass bottom Mattek dishes (MatTek #1.5 coverslips) and transfected with JetPrime reagent according to the manufacturer protocol for GFP-LAMP1. Cells at 37°C were then preincubated for 45min at 25°C to allow the temperature to slowly decrease, then the medium was exchanged in EBSS with 100nM TMR for 6h at 25°C. The medium was exchanged again at time of acquisition in EBSS without phenol red. Movies were recorded on a Leica THUNDER microscope with a Leica HC PL APO 100x/1.44 OIL UV objective. Samples were excited at 475nm and 555nm and collected through 506-532nm and 578-610nm filter cubes. Images were acquired every 9.6s and the temperature of the culture medium was recorded throughout imaging using a semiconductor temperature probe immersed in the medium connected to a CENTER 522 Wireless Datalogger Hygro Thermometer, using the SE522 software (v 3.2.0.0). Deconvolution was performed with the Leica THUNDER Computational Clearing algorithm and image post processing was performed with Adobe Photoshop and ImageJ.

### Correlative electron microscopy (CLEM)

Endogenous tagged eTEX264-Halo were seeded on gridded 35mm glass bottom Mattek dishes (MatTek). Cells were then incubated for 6h in EBSS with 100nM TMR at 25 °C and then fixed for 2h at RT in periodate-lysine-paraformaldehyde (PLP). Samples were then washed in PBS and incubated 10 min with 50 mM glycine and blocked 1h in blocking buffer (0.2% BSA, 5% goat serum, 50mM NH4Cl, 0.1% saponin, 20mM PO4 buffer 150mM NaCl). Cells were stained with primary antibodies, Alexa Fluor-conjugated secondary antibodies and nanogold-labelled secondary antibodies (Nanoprobes) in blocking buffer at RT. Confocal images were acquired with Leica STELLARIS 5 and STELLARIS 8 with a Leica HCX PL APO lambda blue 63x/1.40 oil objective. The software used was Leica LAS X. Cells were then fixed for 30min in 1% glutaraldehyde (GA) and then nanogold particles were enhanced using a gold enhancement solution (Nanoprobes) following the manufacturer’s instructions. Cells were post-fixed with 1% OsO4 and 1.5% potassium ferricyanide in 0.1M cacodylate buffer. Cells were stained with 0.5% uranyl acetate overnight, dehydrated and embedded in Epoxy resin (Sigma-Aldrich) and put in an oven at 60°C for 48h. Ultrathin sections (70-90nm) were cut with an ultramicrotome (UC7, Leica Microsystems), contrasted with uranyl acetate and Sato’s lead solution, and examined with a Talos L120C transmission electron microscope (FEI, Thermo Fisher Scientific) operated at 120 kV. Images were recorded with a Ceta CCD camera (FEI, Thermo Fisher Scientific). For three-dimensional serial-section electron tomography, 130-150nm serial sections were collected on formvar/carbon-coated slot grids, and tilt series (±60°) were acquired on the Talos L120C TEM using Tomography 5 software (FEI, Thermo Fisher Scientific). Tomograms were segmented in Microscopy Image Browser (MIB, v2.91) and visualized in 3dmod (v5.1.3).

### Expansion Microscopy

Cells were seeded on 8 mm coverslips and starved in EBSS for 6h at 25 °C, or at 37 °C in the presence of 100 nM BafA1 where indicated. Samples were cryo-fixed by plunging them in ethane cooled with liquid nitrogen and then freeze substituted. After plunging, coverslips were transferred in 1.5ml vials containing liquid-nitrogen cooled acetone plus 0.5% PFA and 0.02% GA. Vials were incubated in dry ice and slowly brought to 0°C overnight. Rehydration in ethanol was made as follows: ethanol 100%, ethanol 100%, ethanol 95%, ethanol 95%, ethanol 70%, ethanol 50%, ethanol 25% and PBS. Incubations were 5 min for ethanol 100% and 3min for the rest. Expansion microscopy was performed following established protocols(Gambarotto *et al*., 2019; Louvel *et al*., 2023), with minor modifications. Briefly, cryo-fixed coverslips were incubated for 5h in 1.4% PFA / 2% acrylamide (AA) solution in PBS at 37°C. Gelation was then performed by placing coverslips face down on 17µl drops of monomer solution (19% sodium acrylate, 10% AA, 0.1% BIS, 0.25% tetramethylethylenediamine (TEMED)/ammonium persulfate (APS) in 1x PBS) on a parafilm-coated box placed on ice. The first 5 min were done at 4°C, then the box was moved at 37°C. After that, coverslips with gels were incubated in a six-well plate with Denaturation Buffer (DB, 200 mM SDS, 200 mM NaCl, 50 mM Tris-BASE in ddH2O) in agitation until gel detachment, then 1h30min at 95°C in fresh DB. Gels were then placed in ddH2O for expansion, changing water twice after 30 min and leaving it overnight for the last wash. Gels were then incubated for shrinking, washing them twice in PBS for 15 min. Then they were incubated in primary antibodies 1:100 in PBS/2% bovine serum albumin (BSA) for 3h at 37°C under shaking. After that, gels were washed three times for 10 min in PBS/0.1% Tween and then incubated with secondary antibodies 1:300 in PBS/2% BSA for 2h30 min at 37°C under shaking. After that, gels were washed three times for 10 min in PBS/0.1% Tween and placed in ddH2O for expansion, changing water twice after 30 min and leaving it overnight for the last wash. Gels were imaged on a poly-Lysine coated 24mm coverslip with Leica STELLARIS 5 and STELLARIS 8 with a Leica HCX PL APO lambda blue 63x/1.40 oil objective. The software used was Leica LAS X. Deconvolution was performed with Leica Lightning software. Image post-processing was performed with Adobe Photoshop and ImageJ.

### Statistical analysis

Statistical analysis and graphical plots were performed using GraphPad Prism 11 (GraphPad Software, Version 11.1.0). Tests employed for statistical analysis are reported in figure legends. Data distributions (individual points) are always shown. An adjusted *p* value < 0.05 (for one-way analysis of variance (ANOVA) with multiple comparison tests) was considered statistically significant. Results from ANOVA analyses were adjusted for multiple comparisons where appropriate. All t-tests were performed as two-sided tests. The corresponding figure legends indicate the number of cells analyzed (n) and the number of independent biological replicates (N).

## Structured Methods -Reagents and Tools Table

*Instructions: Please complete the relevant fields below, adding rows as needed. The following page provides an example of a completed table and additional instruction for entering your data in the table.*

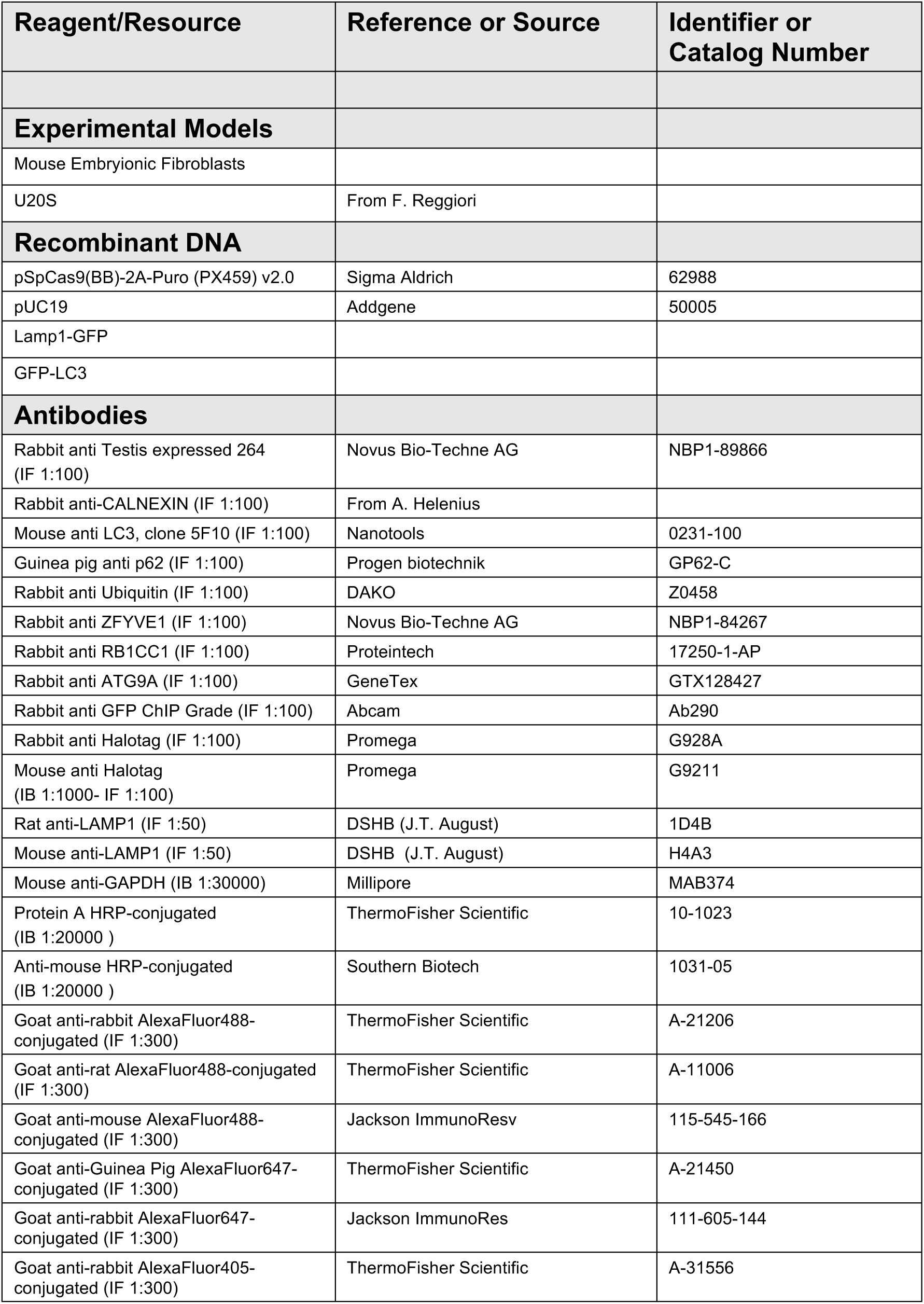
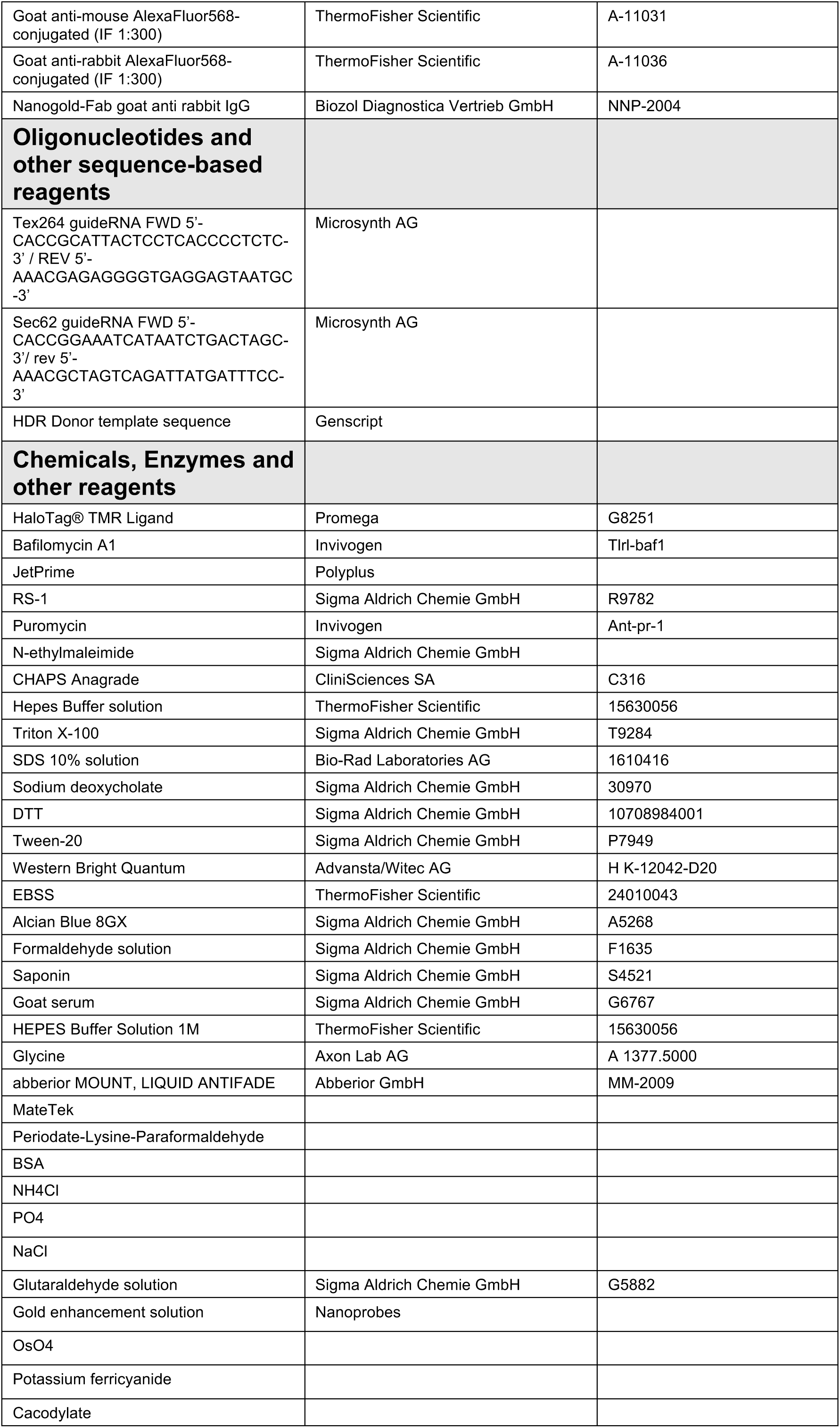
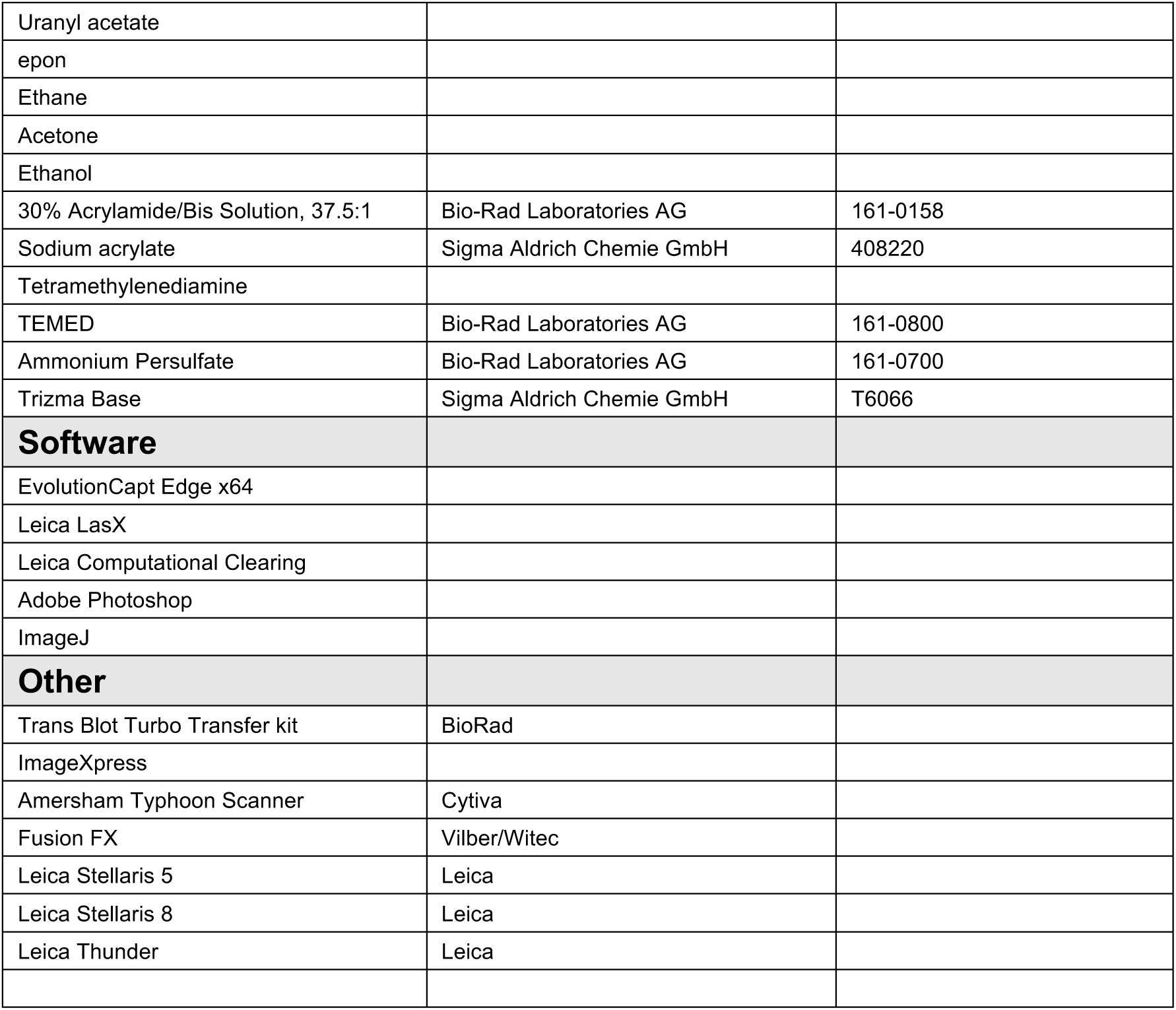

## Acknowledgments

We thank the members of Molinari’s lab for discussions, and critical reading of the manuscript. We thank Fulvio Reggiori for the U2OS cells lacking ATG9A, ATG13, FIP200.

## Funding

Swiss National Science Foundation grants 310030_214903 and 320030-227541 (MM).

## Data availability

This study includes no data deposited in external repositories. Source data for all quantifications are provided with the manuscript. Raw image datasets and tomographic reconstructions are available from the corresponding author on reasonable request.

## Author contributions

MF: Conceptualization; Data curation; Investigation; Visualization; Methodology; Writing-original

draft; Writing-review and editing

TS: Investigation; Visualization; Writing-review and editing

DM: Methodology; Writing—review and editing.

CG: Investigation; Visualization; Writing-review and editing

AR: Methodology; Writing—review and editing.

MM: Conceptualization; Supervision; Funding acquisition; Writing-original draft; Project administration; Writing-review and editing.

## Disclosure and competing interests statement

The authors declare that they have no conflict of interest.

